# Human synaptic neoteny requires species-specific balancing of SRGAP2-SYNGAP1 cross-inhibition

**DOI:** 10.1101/2023.03.01.530630

**Authors:** Baptiste Libé-Philippot, Ryohei Iwata, Aleksandra J Recupero, Keimpe Wierda, Martyna Ditkowska, Vaiva Gaspariunaite, Ben Vermaercke, Eugénie Peze-Heidsieck, Daan Remans, Cécile Charrier, Franck Polleux, Pierre Vanderhaeghen

## Abstract

Human-specific (HS) genes are potential drivers of brain evolution, but their impact on human neuron development and disease remains unclear. Here we studied HS genes SRGAP2B/C in human cortical projection neurons (CPNs) in vivo, using xenotransplantation in the mouse cortex. Downregulation of SRGAP2B/C in human CPNs greatly accelerated synaptic development, indicating their requirement for human-specific synaptic neoteny. SRGAP2B/C acted by downregulating their ancestral paralog SRGAP2A, thereby upregulating postsynaptic levels of SYNGAP1, a major intellectual deficiency/autism spectrum disorder (ID/ASD) gene. Combinatorial genetic invalidation revealed that the tempo of synaptogenesis is set by a balance between SRGAP2A and SYNGAP1, which in human CPNs is tipped towards neoteny by SRGAP2B/C. Our results demonstrate that HS genes can modify the phenotypic expression of ID/ASD mutations through regulation of synaptic neoteny.

**One-Sentence Summary:** Human-specific genes SRGAP2B/C control human cortical neuron neoteny by regulating the function of neurodevelopmental disorder gene SYNGAP1

## Main Text

A salient feature of human brain development is the considerably prolonged tempo of maturation of cortical neurons, taking months to years in humans, instead of days to weeks in other mammals (*1–4*). Cortical neuron neoteny is thought to lead to enhanced synaptic plasticity and cognitive functions characterizing *Homo Sapiens* (*2, 5–7*). Conversely, accelerated brain development has been associated with some neurodevelopmental diseases (NDD) that impair higher cognitive functions, leading to intellectual deficiency (ID) and autism spectrum disorder (ASD) (*8*). The mechanisms by which NDDs affect human-specific circuit development and function remain unclear: they could affect molecular pathways that uniquely control neoteny in our species, but this remains to be tested.

Despite the significance of cortical neuron neoteny for human brain evolution and (dys)function, the underlying molecular and cellular mechanisms remain poorly understood. Human cortical projection neurons (CPNs) develop at their species-specific tempo when xenotransplanted in the fast-developing mouse cortex, pointing to cell-intrinsic mechanisms (*9–13*). While mitochondria metabolism was recently shown to regulate the species-specific tempo of global neuronal maturation (*14*), one outstanding molecular candidate for the specific timing of synaptic maturation is the SRGAP2 gene family (*15*). This family is composed of SRGAP2A, the ancestral gene shared by all mammals, and human-specific (HS) gene duplicates SRGAP2B, SRGAP2C and SRGAP2D, which emerged through segmental duplications during recent hominin evolution (*16, 17*). Studies in mouse revealed that SRGAP2A is a postsynaptic protein that acts as a positive regulator of CPN synaptic maturation (*15, 18*), while the HS paralogs SRGAP2B and SRGAP2C encode a truncated version of the F-BAR domain found in SRGAP2A (*15*). Expression SRGAP2B or SRGAP2C in developing mouse CPNs, or downregulation of SRGAP2A, lead to delayed maturation of dendritic spines and synapse development with a higher final synaptic density in adult cortical neurons, mimicking some aspects of human neuronal neoteny and adult connectivity (*15, 18, 19*). While these data point to the SRGAP2 genes as important regulatory components of synaptic development and evolution, it remains unclear if and how HS genes SRGAP2B/C contribute to human neuron neoteny, whether in physiological or pathological conditions.

### An in vivo model of SRGAP2 loss of function in human CPNs

Here we explored the function and mechanisms of action of SRGAP2B/C in human CPNs vivo, using xenotransplantation in the mouse neonatal cortex. In order to downregulate in human CPNs both SRGAP2B/C simultaneously without affecting SRGAP2A expression, we used a knock- down (KD) approach with lentiviral vectors expressing two independent shRNAs targeting specifically the 3’-untranslated region (3’-UTR) of SRGAP2B and SRGAP2C transcripts, which is not present in the ancestral SRGAP2A mRNA transcript (Fig.1A) (*15*). This approach was validated in human CPNs differentiated from pluripotent stem cells (PSC) for 45 days in vitro, leading to reduction of SRGAP2B/C proteins (Fig. S1A). Conversely, to examine the role of the ancestral SRGAP2A gene in human neurons, we used a SRGAP2A KD approach, which led to decreased SRGAP2A protein expression in human CPNs (Fig. S1A).

**Fig. 1.**
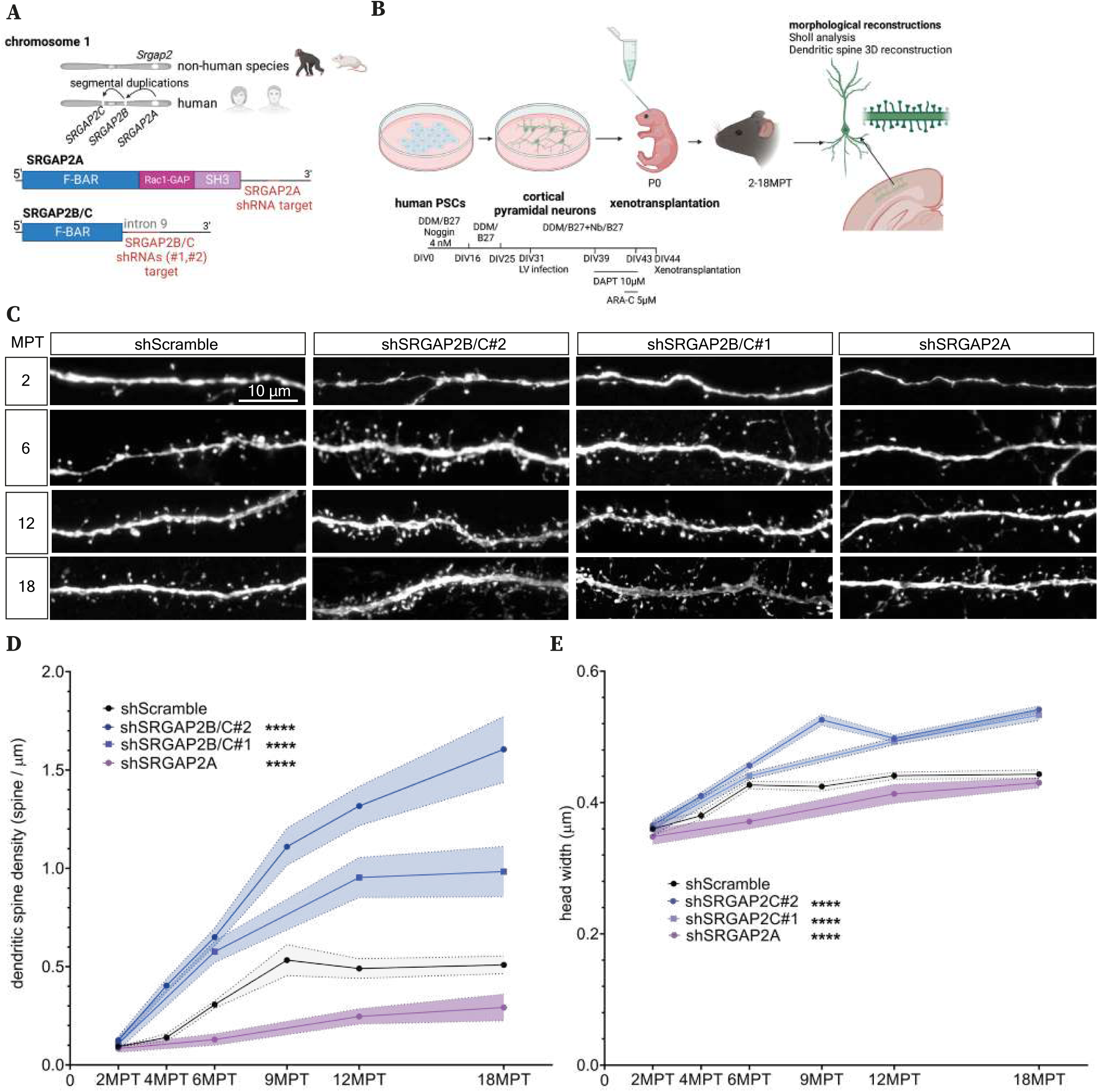
SRGAP2B/C genes are required for dendritic spine neoteny of human cortical pyramidal neurons. (**A**) Description of the SRGAP2 gene family at the genomic and transcript levels; note the location of the mRNA sequences targeted by shRNAs and that SRGAP2B/C shRNAs do not target the SRGAP2C protein coding sequence (cDNA, in blue). (**B**) Experimental design of human pluripotent stem cell (PSC)-derived neurons, infected in vitro with lentivirus (LV) expressing EGFP and shRNAs targeting Scramble, SRGAP2B/C or SRGAP2A sequences +/- SRGAP2C-HA cDNA, and then xenotransplanted in the mouse neonatal cerebral cortex followed by morphological analyses (see Methods). (**C**) Representative proximal dendritic branches of human PSC-derived neurons xenotransplanted in the mouse cerebral cortex, with LV-shRNA infection, followed from 2 month-post-transplantation (MPT) to 18MPT. (**D**) Corresponding quantifications of the dendritic spine density (means +/- SEM; 2-way ANOVA tests)(numbers in Table S1; 9-72 neurons, 2-14 animals, 1-8 litters, per stage per condition). (**E**) Corresponding quantifications of the dendritic spine head width (means +/- 95% confident interval; 2-way ANOVA tests; numbers in Table S1; 9-72 neurons, 101-2007 dendritic spines, 2-14 animals, 1-8 litters, per stage per condition). ****, p<0.0001.

To assess the role of SRGAP2B/C in differentiation and synaptic development of human CPNs in vivo, we used a model of xenotransplantation in the neonatal mouse cerebral cortex (*9, 11*) (Fig. 1B). Using this experimental approach, human cortical progenitors derived from PSC are infected in vitro with lentiviral vectors transducing the shRNAs, as well as EGFP to enable identification and morphological analysis of the transplanted human neurons (Fig. S1B). Shortly before transplantation the human cortical cells are treated with a γ-secretase inhibitor (DAPT) and antimitotic drug (Cytarabine, ARA-C) in order to enhance neurogenesis and enrich for cohorts of transplanted neurons that became postmitotic synchronously (*11*). Following transplantation into the host mouse at P0 (newborn mouse), human CPNs of similar birthdate integrate morphologically and functionally in the mouse cerebral cortex and mature following their species-specific timeline, taking months to develop complex dendritic morphology, mature dendritic spines, functional synapses, and integration into the host cortical circuits (*11*).

### SRGAP2B/C are required for morphological neoteny in human CPNs in vivo

We first analyzed the transplanted human CPNs at an early stage (2 months post-transplantation (MPT)) to assess their integration and cell fate. SRGAP2B/C KD and control neurons integrated similarly in the mouse cortical gray matter, and expressed mostly deep-layer 5/6 markers in similar proportions, as previously described (Fig. S1B-C) (*11*). These data indicate that SRGAP2B/C downregulation had no detectable impact on cortical neuron integration and fate determination, in line with the data obtained in the mouse following expression of SRGAP2C (*15, 20*). We next analyzed the morphological development of the xenotransplanted human CPNs at 2 and 6 MPT, a period during which cortical neurons develop increasingly complex dendritic arborization (*11*). Surprisingly, this revealed that SRGAP2B/C KD at 2MPT displayed a higher degree of dendritic complexity, quantified using Sholl analysis, compared with human CPNs transduced with control (scrambled) shRNA (Fig. S2A-B), while this was not observed at 6MPT (Fig. S2E-F). SRGAP2A KD neurons did not reveal differences in their cortical integration, fate marker expression and dendritic arbor complexity at any stage examined (Fig. S2A-B,E-F).

To assess the specificity of the phenotypes observed, we performed a rescue experiment by co-infecting the SRGAP2B/C KD neurons with a lentivirus expressing the SRGAP2C coding cDNA, which is not targeted by the shRNAs, prior to xenotransplantation. This was sufficient at 2MPT to fully abolish the effects of the SRGAP2B/C KD, confirming the specificity of our shRNAs targeting human-specific paralogs SRGAP2B/C (Fig. S2C-D). These results suggest that HS genes SRGAP2B/C transiently contribute to neotenic dendritic maturation.

We next analyzed dendritic spine maturation, the main cellular target of SRGAP2A downregulation or SRGAP2C expression in mouse CPNs, in the transplanted human CPNs along an extended developmental timeline (from 2 to 18 MPT) (*15*).

At 2MPT, spine density (an index of synaptogenesis) and spine head size (an index of synaptic maturation) were low and similar between all groups (Fig. 1C-E). However, from 4MPT to 18MPT, SRGAP2B/C KD neurons displayed a striking acceleration in the rate of dendritic spine formation (Fig. 1C-D) as well as increased spine head width compared with control human CPNs (Fig. 1C,E). Conversely, in SRGAP2A KD neurons, we observed a slower rate of spine formation (reduced spine density over time; Fig. 1C-D) and a decrease in spine head width (Fig. 1C,E).

At 18MPT, SRGAP2B/C KD human CPNs displayed a 2-3 fold increase in spine density compared to control human CPNs (Fig. 1D). In comparison, the dendritic spine density of deep-layer CPNs of the human prefrontal cortex at one year postnatally is around 0.3-0.5 spines per µm, similarly to our control values, and reaches values of around ∼1 spine per µm around 5-10 years (y) postnatally (*21*). Thus, SRGAP2B/C-deficient human neurons display a strong acceleration of spine formation and maturation.

We next performed rescue experiments as above using SRGAP2C cDNA combined with SRGAP2B/C shRNAs, which was sufficient at 6MPT to abolish the increased density and head width of the dendritic spines induced by SRGAP2C KD (in Fig. S2H-J, differences between shScramble or shScramble+SRGAP2C and shSRGAP2B/C+SRGAP2C are not significant, p>0.05), confirming the specificity of the KD approach (Fig. S2G-J).

These results indicate that HS genes SRGAP2B/C are necessary for the neotenic formation and maturation of dendritic spines in human CPNs in vivo, while downregulation of SRGAP2A leads to decreased pace of dendritic spine formation and maturation.

### SRGAP2B/C are required for functional neoteny in human CPNs in vivo

Next, we determined the functional consequences of the accelerated development of dendritic spines of SRGAP2B/C KD neurons using patch clamp recordings of human CPNs in *ex vivo* cortical slices prepared from xenotransplanted mice (Fig. 2A). We focused on spontaneous excitatory post-synaptic currents (sEPSC), which typically increase in frequency and amplitude in xenotransplanted human CPNs over the first 6 MPT, reflecting increasing synapse formation and maturation, respectively (*11*). We observed a significant increase of sEPSC frequency in SRGAP2B/C KD neurons compared to control neurons (Fig. 2B-C), indicating increased number of functional synapses (*22*) as well as a significant increase in sEPSC amplitude in SRGAP2B/C KD neurons compared to control neurons (Fig. 2B,D), suggesting accelerated rates of excitatory synaptic maturation (*22*).

**Fig. 2.**
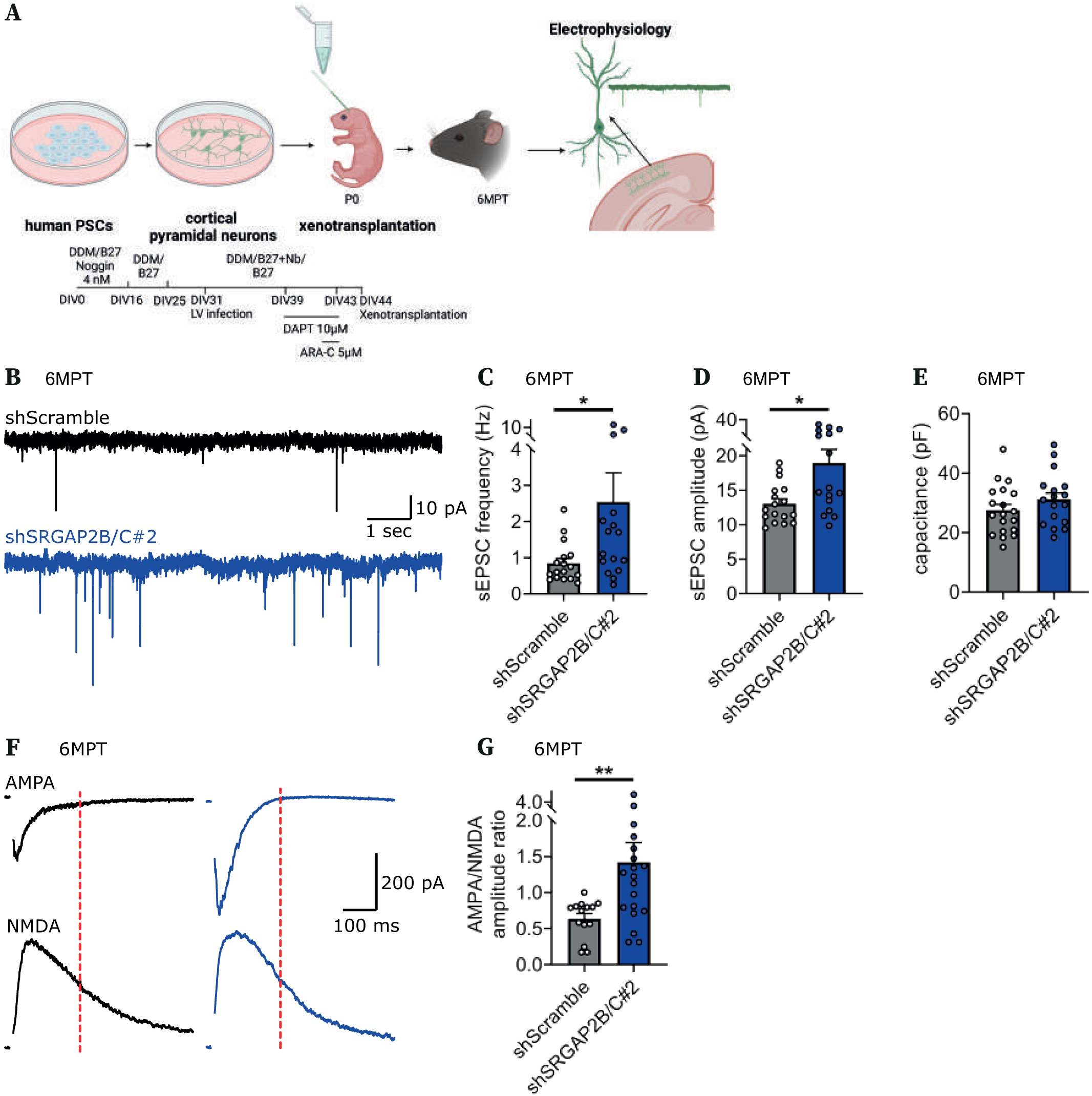
SRGAP2B/C are required for neotenic synaptic maturation of human cortical neurons. (**A**) Experimental design of human PSC-derived neurons, LV infected in vitro expressing EGFP and shRNAs targeting Scramble or SRGAP2B/C sequences, and then xenotransplanted in the mouse neonatal cerebral cortex followed by electrophysiological analyses at 6MPT. (**B**) Representative traces of synaptic currents of human PSC-derived neurons xenotransplanted in the mouse cerebral cortex, with LV-shRNA infection, at 6MPT. (**C-D**) Corresponding quantification of the sEPSCs frequency and amplitude (Mann-Whitney tests)(16-17 neurons per condition from 6-7 animals from 2 litters). (**E**) Cell capacitance of human neurons, with LV-shRNA infection, at 6MPT (Mann-Whitney test)(16-17 neurons per condition from 6-7 animals from 2 litters). (**F**) Representative examples of AMPA and NMDA currents of human neurons, with LV-shRNA infection, at 6MPT. (**G**) Corresponding quantification of the AMPA/NMDA currents amplitude ratio (Mann-Whitney tests)(14-19 neurons per condition from 5-7 animals from 2 litters). Data are represented as mean + SEM. *, p<0.05; **, p<0.01.

Under voltage clamp conditions, SRGAP2B/C KD neurons did not differ from control neurons regarding their (measurable) membrane capacitance (Fig. 2E), consistent with similar dendritic arbor size at this stage (Fig. S2E-F). We next measured AMPA and NMDA currents as AMPA/NMDA current ratios represent a well-validated index of postsynaptic maturation (*22*). This revealed a >2-fold increase in AMPA/NMDA ratios in SRGAP2B/C KD neurons compared to control neurons (Fig. 2F-G), confirming that postsynaptic maturation reaches much higher levels following SRGAP2B/C loss of function.

These results demonstrate that SRGAP2B/C-deficient human CPNs display functionally more mature and stronger excitatory synapses, containing higher AMPA/NMDA receptor ratios than in control human CPNs, revealing the requirement of SRGAP2B/C for functional synaptic neoteny in human CPNs.

### SRGAP2B/C regulates the synaptic levels of SRGAP2A and SYNGAP1

Given the strong effects of SRGAP2B/C loss of function on the timing of synaptic development of human neurons in vivo, we explored the underlying molecular mechanisms. Work in mouse CPNs suggests that SRGAP2B/C acts by antagonizing the function of its ancestral gene SRGAP2A (*15, 18*). Studies in heterologous expression systems indicate that SRGAP2B/C can heterodimerize with SRGAP2A, leading to the formation of insoluble complexes that are degraded by the proteasome (*19, 23*). We therefore examined SRGAP2A protein levels in human CPNs in cytosolic and synaptosomal fractions following SRGA2B/C KD, SRGA2A KD, or control human CPNs, infected at DIV45 and cultured in vitro until DIV70 (Fig. 3A). As predicted by previous results obtained in mouse CPNs, SRGAP2B/C KD in human CPNs led to significant increase of SRGAP2A protein levels in the cytosolic fraction and most significantly in the synaptosome compartment (Fig. 3B-C, Fig. S3A-B), demonstrating that human-specific paralogs SRGAP2B/C negatively regulate the levels of SRGAP2A in human CPNs and synapses.

**Fig. 3.**
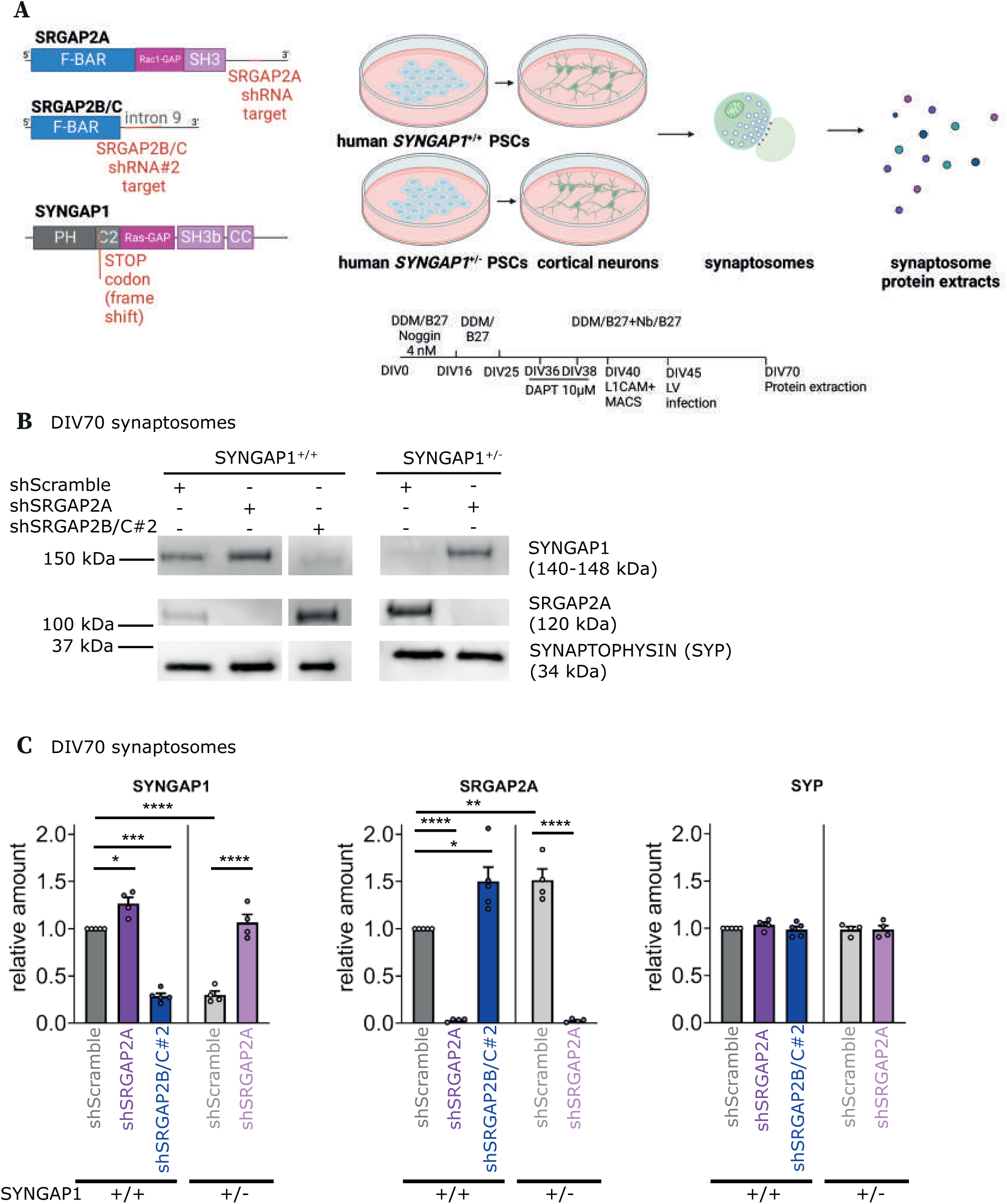
SRGAP2B/C are human-specific regulators of SRGAP2A/SYNGAP1 antagonism at the synapse. (**A**) Experimental design: synaptosomal protein extraction at day in vitro (DIV)70 from human SYNGAP1^+/+^ and SYNGAP1^+/-^ PSC-derived neurons with LV-shRNA infection at DIV45. (**B**) Synaptic fraction of SYNGAP1^+/+^ and SYNGAP1^+/-^ PSC-derived neurons at DIV70 with LV-shRNA infection at DIV45 stained for SYNGAP1, SRGAP2A and SYP. (**C**) Corresponding quantifications (ANOVA multiple tests; 3-5 experiments per condition). Data are represented as mean + SEM. *, p<0.05; **, p<0.01; ***, p<0.001; ****, p<0,0001).

Given the increase in SRGAP2A protein abundance at the synapse following SRGAP2B/C KD, we probed for potential changes in other synaptic proteins. We first examined Homer1 and PSD95, two key scaffolding proteins of AMPA and NMDA receptors at excitatory synapses but found no detectable protein level changes following SRGAP2B/C, or SRGAP2A KD at these early stages of neuronal development (Fig. S3C-D).

We next turned to SYNGAP1, an abundant post-synaptic protein that acts as a negative regulator of excitatory synaptic maturation (*24, 25*) and is encoded by a gene frequently mutated in ID/ASD (*26–28*). Indeed, using the same model of xenotransplantation, we recently found that human CPNs displaying SYNGAP1 pathogenic mutations display a striking acceleration in dendritic spine formation and synaptic maturation (*29*), highly reminiscent what we observed here following SRGAP2B/C KD. Remarkably, we found that SYNGAP1 protein levels are dramatically decreased in the synaptosome fraction of SRGAP2B/C KD human CPNs, while its cytosolic levels remained unchanged (Fig. 3B-C, Fig. S3A-B). Conversely, we found that SYNGAP1 protein levels were significantly increased in synaptosomes isolated from SRGAP2A KD neurons, while no difference was detected in the cytosolic fraction compared to control neurons (Fig. 3B-C, Fig. S3A-B). These results indicate that SRGAP2A inhibits the synaptic accumulation of SYNGAP1. We next examined whether the converse is the case, i.e. whether SYNGAP1 negatively regulates the synaptic levels of SRGAP2A, by examining their protein levels in human CPNs generated from SYNGAP1 haploinsufficient (SYNGAP1^+/-^) PSC (generated in (*29*)). As expected, SYNGAP1^+/-^ human CPNs displayed a 50% reduction in SYNGAP1 levels in cytosolic compartments (Fig. S3A-B) and >50% in synaptic compartments (Fig. 3B-C). Strikingly, we also detected significantly increased synaptic levels of SRGAP2A in SYNGAP1^+/-^ human CPNs compared to control SYNGAP1^+/+^ human CPNs (Fig. 3B-C), while no changes were detected in the cytosolic fraction (Fig. S3A-B). SRGAP2B/C levels on the other hand did not display any detectable changes in SYNGAP1^+/-^ neurons compared to control human CPNs (Fig. S3A-B). Finally, we found that SRGAP2A KD in SYNGAP1^+/-^ human neurons led to an increase in SYNGAP1 levels at the synapse, reaching levels comparable to those found in SYNGAP1^+/+^ control human neurons (Fig. 3B-C), while no change was detected in the cytosolic fraction (Fig. S3A-B).

Overall, these results indicate that SRGAP2A and SYNGAP1 cross-inhibit their synaptic accumulation in developing human cortical neurons, and that HS genes SRGAP2B/C, by downregulating SRGAP2A, increase the synaptic levels of SYNGAP1 in human CPNs (Fig. 4L).

**Fig. 4.**
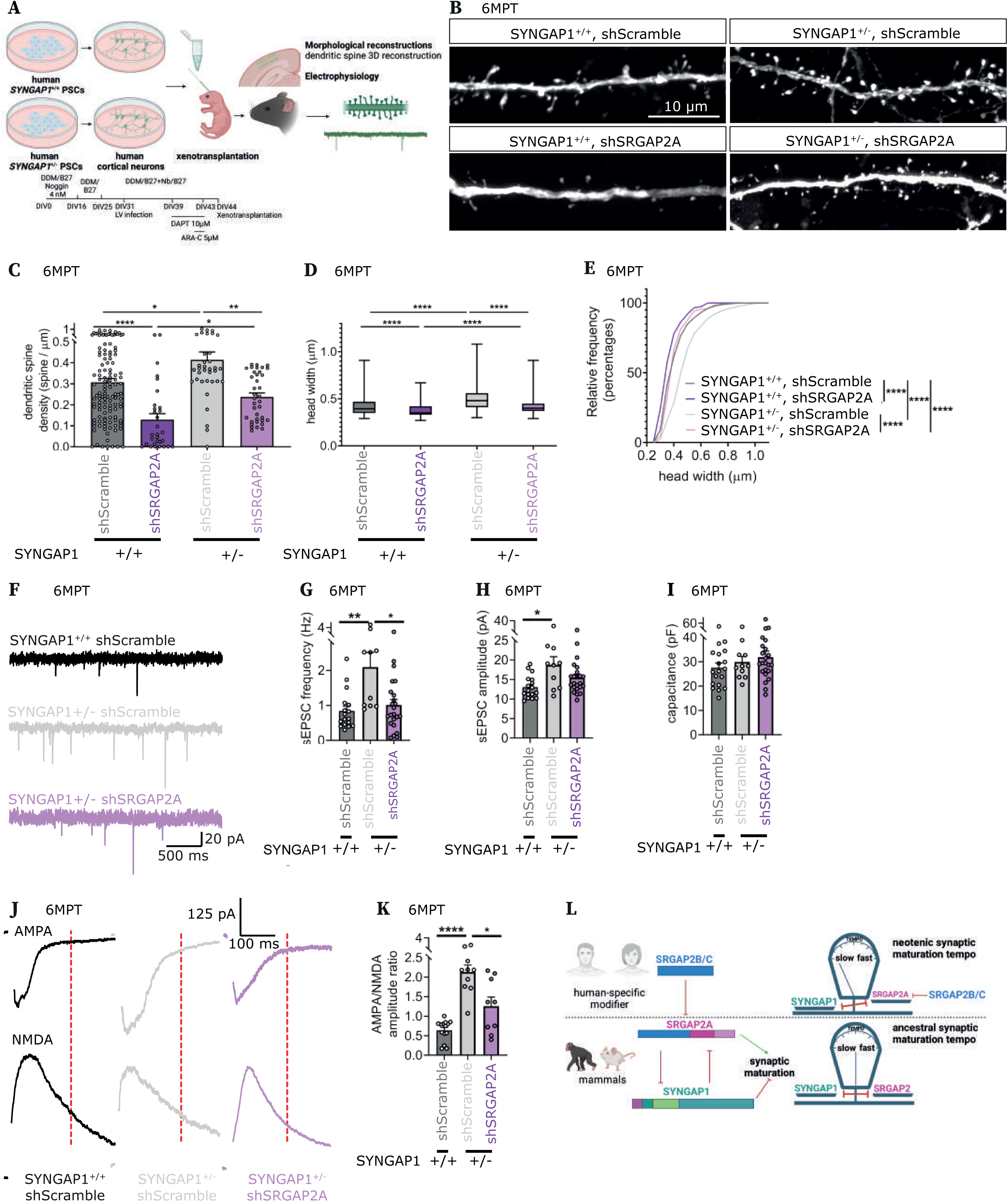
SRGAP2A and SYNGAP1 epistasis regulates neuronal developmental tempo in human. (**A**) Experimental design of SYNGAP1^+/+^ and SYNGAP1^+/-^ human PSC-derived neurons, LV infected in vitro expressing EGFP and shRNAs targeting Scramble and SRGAP2A sequences, and then xenotransplanted in the mouse neonatal cerebral cortex followed by morphological and electrophysiological analyses at 6MPT. (**B**) Representative proximal dendritic branches of SYNGAP1^+/+^ and SYNGAP1^+/-^ human PSC-derived neurons xenotransplanted in the mouse cerebral cortex, with LV-shRNA infection, at 6MPT. (**C-D**) Corresponding quantification of the dendritic spine density and spine head width (Kruskal-Wallis tests)(for SYNGAP1^+/+^ neurons: 15- 72 neurons, 268-1440 dendritic spines from 5-14 animals from 2-8 litters per condition; for SYNGAP1^+/-^ neurons: 19 neurons, 689-908 dendritic spines from 6 animals from 2 litters per condition)(SYNGAP1^+/+^ data were already shown in Fig. 1D-E). (**E**) Corresponding cumulative distribution of the spine head width (Kolmogorov-Smirnov tests). (**F)** Representative examples of synaptic currents of SYNGAP1^+/+^ and SYNGAP1^+/-^ human PSC-derived neurons xenotransplanted in the mouse cerebral cortex, with LV-shRNA infection, at 6MPT. (**G-H**) Corresponding quantifications of the sEPSCs frequency and amplitude (Kruskal-Wallis tests)(for SYNGAP1^+/+^ neurons: 18 neurons from 6 animals from 2 litters; for SYNGAP1^+/-^ neurons: 10-23 neurons from 5-6 animals from 3 litters per condition)(SYNGAP1^+/+^ shScramble data were already shown in Fig. 2C-D). (**I**) Cell capacitance of SYNGAP1^+/+^ and SYNGAP1^+/-^ human PSC-derived neurons xenotransplanted in the mouse cerebral cortex, with LV-shRNA infection, at 6MPT (Kruskal-Wallis tests)(for SYNGAP1^+/+^ neurons: 18 neurons from 6 animals from 2 litters; for SYNGAP1^+/-^ neurons: 10-23 neurons from 5-6 animals from 3 litters per condition)(SYNGAP1^+/+^ shScramble data were already shown in Fig. 2E). (**J)** Representative examples of AMPA and NMDA currents of SYNGAP1^+/+^ and SYNGAP1^+/-^ human PSC-derived neurons xenotransplanted in the mouse cerebral cortex, with LV-shRNA infection, at 6MPT. (**K**) Corresponding quantification of the AMPA/NMDA currents amplitude ratio (Kruskal-Wallis tests)(for SYNGAP1^+/+^ neurons: 15 neurons from 5 animals from 2 litters; for SYNGAP1^+/-^ neurons: 10 neurons from 3 animals from 3 litters per condition)(SYNGAP1^+/+^ shScramble data were already shown in Fig. 2G). (**L**) Schematic representation of the main findings: SRGAP2A and SYNGAP1 have an antagonistic function of the synaptic maturation of cortical pyramidal neurons in mammals which is regulated in a species-specific fashion in human by SRGAP2B/C. In C, G-I, K, data are represented as mean+SEM. *, p<0.05; **, p<0.01; ****, p<0.0001.

### The balance between SYNGAP1 and SRGAP2A/B/C determines human synaptic neoteny

As SYNGAP1 slows down the rate of excitatory synapse maturation (*24, 29*) while SRGAP2A exerts the opposite effect (Fig. 1C-E) (*15*), we next tested if the molecular antagonism we uncovered had an impact on the rate of synaptic maturation in human CPNs in vivo. Specifically, we tested whether the accelerated rate of synaptic maturation characterizing SYNGAP1^+/-^ human CPNs compared to control human neurons (*29*) could be rescued by interference with SRGAP2A. To test this, we performed xenotransplantation of SYNGAP1^+/-^ human CPNs, infected with control or SRGAP2A KD shRNAs, followed by morphological and functional synaptic characterization at 6MPT compared to control (SYNGAP1^+/+^) neurons (Fig. 4A).

We first focused on dendritic spine maturation. As shown above, SRGAP2A KD led to a significant decrease in dendritic spine density and spine head width (Fig. 1C-E, Fig. 4B-E). Conversely, SYNGAP1^+/-^ human CPNs displayed increased dendritic spine density, as we previously observed (*29*)) and larger head width (Fig. 4B-E). Most remarkably, we found that the combined loss of function of SRGAP2A and SYNGAP1 led to normalization of both dendritic spine density and spine head size in human CPNs at 6MPT (Fig. 4B-E), while differences between SYNGAP1^+/+^ shScramble and SYNGAP1^+/-^ shSRGAP2A are not significant (p>0.05; Fig. 4C-D). We next assessed functional synaptic maturation of human CPNs following the same manipulations using patch clamp recordings of xenotransplanted human neurons in *ex vivo* cortical slices isolated at 6MPT. We found that SYNGAP1^+/-^ human CPNs displayed increased sEPSC frequency and amplitude, as well as increased AMPA/NMDA ratio, consistent with increased rates of synaptic maturation compared to control SYNGAP1^+/+^ human neurons (Fig. 4F-K as previously found in (*29*)). Strikingly, these phenotypes were rescued following SRGAP2A KD (Fig. 4F-K), while differences between SYNGAP1^+/+^ shScramble and SYNGAP1^+/-^ shSRGAP2A are not significant (p>0.05; Fig. 4G-H,K), further indicating strong antagonism between SYNGAP1 and SRGAP2A regarding their effects on functional synaptic maturation in human CPNs.

These results indicate that the tempo of synapse development in human CPNs is regulated by the antagonistic balance between SRGAP2A and SYNGAP1, which is tipped by SRGAP2B/C, leading to neoteny of synaptic maturation (Fig. 4L).

### SRGAP2/SYNGAP1 antagonism during synaptic maturation is conserved in mice

Is the SRGAP2/SYNGAP1 antagonism in the control of dendritic spine maturation shared with other mammalian species lacking SRGAP2B/C, such as the mouse? We reasoned that if ancestral SRGAP2 promotes dendritic spine maturation at least in part by limiting or antagonizing SYNGAP1 accumulation postsynaptically, then the delayed synaptic maturation observed in mouse CPNs heterozygous for the SRGAP2A mouse orthologous gene Srgap2 (*15, 18*) should be suppressed in Srgap2/Syngap1 double heterozygous neurons. To test this, we crossed two conditional knockout mouse lines for Srgap2 and Syngap1 and induced sparse, Cre-recombinase dependent, cell autonomous haploinsufficiency of Srgap2, Syngap1, or both, in layer 2/3 CPNs (Fig. 5A-B). As reported previously (*15, 18, 24*), analysis at P21 revealed that dendritic spine head size was significantly smaller in Srgap2 heterozygous CPNs, and significantly larger in Syngap1 heterozygous CPNs compared to WT layer 2/3 CPNs (Fig. 5C-E). Most interestingly, we observed an almost complete normalization of dendritic spine head size distribution in Srgap2^F/+^;Syngap1^F/+^ double heterozygotes CPNs bringing the values very close to WT CPNs (Fig. 5C-E). In line with our results in human neurons (Fig. 4B-E), these results demonstrate that SRGAP2 and SYNGAP1 cross-inhibit each other in dendritic spine maturation in mouse CPNs in vivo.

**Fig. 5.**
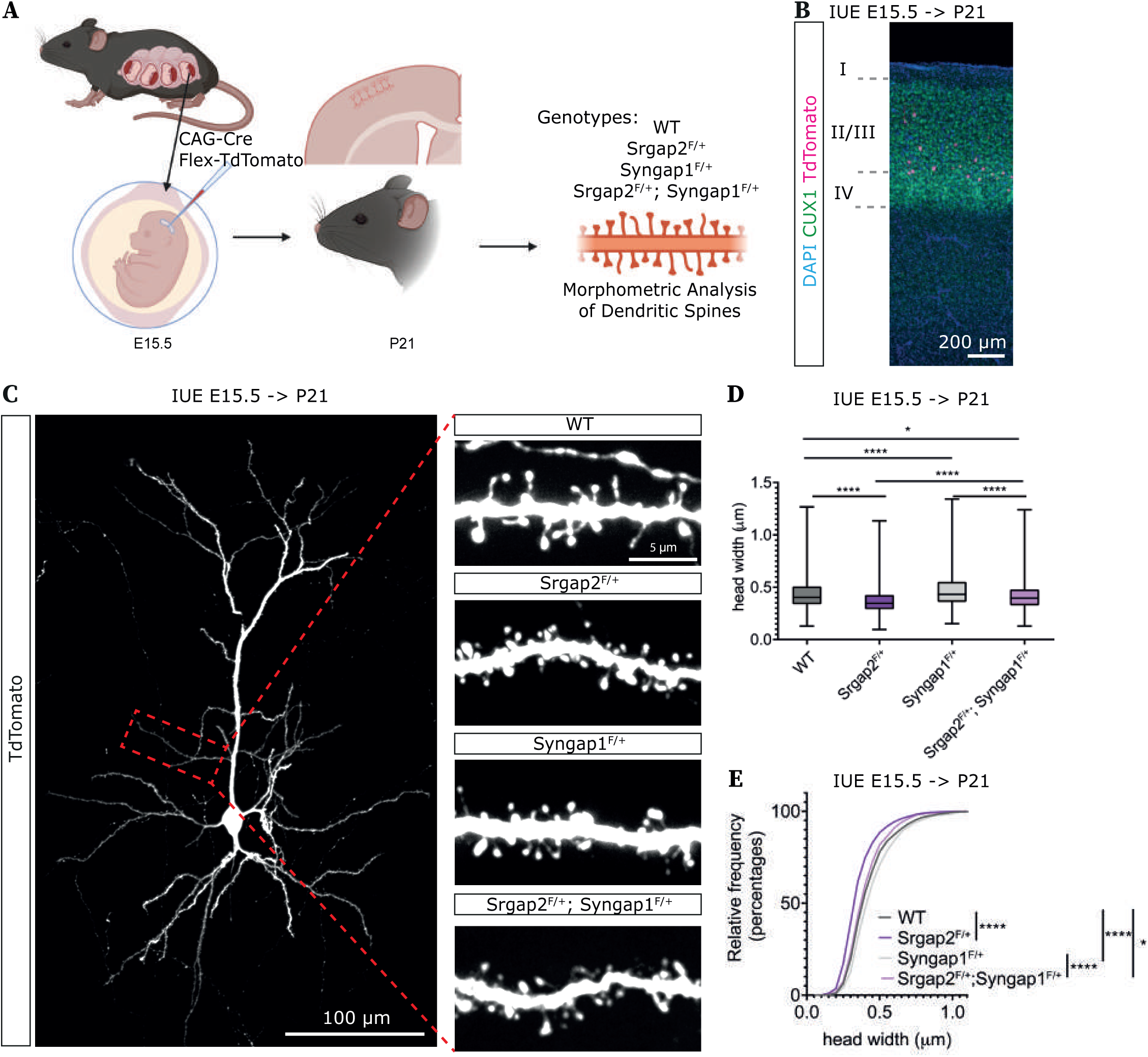
SRGAP2A and SYNGAP1 epistasis regulates neuronal developmental tempo in mouse cortex. (A) Experimental design of in utero electroporation (IUE) at embryonic day 15.5 (E15.5) that targets the flex-TdTomato construct and cre recombinase to layer 2/3 CPNs of mouse embryos. Morphometric dendritic spine analysis was then performed on postnatal day (P) 21 coronal sections of the following genotypes: WT, Srgap2^F/+^, Syngap1^F/+^ and Srgap2^F/+^;Syngap1^F/+^. (**B**)Representative WT coronal section following IUE of the mouse barrel cortex and immunostained for CUX1 (magenta) and DAPI (blue). (**C**) Representative optically isolated L2/3 CPN from P21 WT mouse. Insets represent apical oblique dendritic segments of the respective genotype. (**D**) Corresponding quantification of the spine head widths (Kruskal-Wallis tests)(18-21 neurons, 1030-1345 dendritic spines, 3-4 mice from 2-3 litters per condition). (**E**) Corresponding cumulative distribution of the dendritic spine head widths (Kolmogorov-Smirnov tests). *, p<0.05; ****, p<0.0001.

## Discussion

Cortical neuron neoteny is thought to play major role in human-specific brain function and plasticity, but the underlying molecular mechanisms remain poorly understood (*3, 4, 30*). Here we demonstrate that HS genes SRGAP2B/C are necessary for the neotenic synaptic development and maturation characterizing human CPNs. We identified an antagonistic interaction between postsynaptic proteins SRGAP2A and SYNGAP1 that sets the tempo of synaptic maturation of mammalian pyramidal neurons and found that this balance is modulated in a species-specific manner by SRGAP2B/C, which ‘tips the balance’ towards SYNGAP1, thereby setting a neotenic tempo of synaptic maturation in human CPNs.

Our finding that SRGAP2B/C KD leads to increased spine density in human CPNs may appear paradoxical given the positive effects of inducing SRGAP2C on synaptic density in mature mouse CPNs (*15, 18, 20*). This likely reflects the fact that our in vivo xenotransplantation model, even after 18 MPT, corresponds to relatively early stages of human CPN development, during which spine density reflects the rates of synapse formation. Allowing differentiation of human CPNs for significantly longer periods of time could reveal additional roles for SRGAP2C in regulating the final number of synapses in more mature human neurons. However, we are currently limited by the life span of host species such as the mouse.

At the molecular level, we found that downregulation of SRGAP2B/C increases SRGAP2A synaptic protein levels in human CPNs, as previously suggested in heterologous systems (*19, 23*). Most strikingly, our study uncovers a molecular antagonism between SRGAP2A and SYNGAP1, two synaptic proteins that exert opposite roles in developmental timing of synapse formation and maturation. SYNGAP1 is key regulator of synaptic formation and function in cortical neurons (*25*) and *SYNGAP1* haploinsufficiency is responsible for 1% of all cases of non-syndromic ID, frequently associated with ASD (*25, 31*). Previous work has pointed to precocious synapse formation in Syngap1^+/-^ juvenile mutant mice (*24*). Using xenotransplantation, SYNGAP1 haploinsufficiency in human CPNs in vivo was recently found to lead to strong acceleration of synaptogenesis and synaptic maturation (*29*). This phenotype is strikingly similar to what we observed here following SRGAP2B/C loss of function, consistent with our finding that HS genes SRGAP2B/C promotes neoteny by upregulating the synaptic levels of SYNGAP1 in human CPNs. It remains to be determined whether and how direct or indirect interactions between SRGAP2A and SYNGAP1 proteins mediate their cross-inhibition at the excitatory synapse. SYNGAP1 and SRGAP2A could compete for the same postsynaptic binding partner, yet to be determined in a mutually exclusive fashion. In addition or alternatively, CTNND2, which was recently found to interact with both SRGAP2A and SYNGAP1 (*32*) could be a pivotal postsynaptic protein regulating SRGAP2A/SYNGAP1 antagonism.

Precocious brain development was previously associated with specific forms of NDD including ASD (*33, 34*), and in vitro PSC models have recently suggested altered timing of neuronal development in some forms of ASD (*35*). Our findings strongly suggest that alterations of the neotenic pace of synaptic maturation in human CPNs may be a critical pathogenic mechanism in some forms of NDD, which will be important to further explore clinically and experimentally. Our data indicate that human CPNs are particularly sensitive to SRGAP2B/C levels of expression: after 18 months, the SRGAP2B/C KD CPNs display levels of synaptic maturation observed for human CPNs after 5 years in vivo (*21*). This suggests that alterations of SRGAPB/C levels or function could have pathological consequences in the human brain in vivo, potentially ASD and/or ID symptoms. While no evidence for SRGAP2B/C germline mutations or copy-number variations (CNV) have been reported yet, the possibility of rare pathogenic variants remains to be explored (*36–38*). Finally, by linking HS genes to SYNGAP1, a gene frequently mutated in ID/ASD, our study identifies molecular mechanisms by which the phenotypic expression of genetic mutations leading to neurodevelopmental disorders can be human-specific.

## Acknowledgments

We thank members of the PV lab for helpful discussions and precious help. We thank Joris de Wit lab (CBD) for excellent advice and Bart de Strooper lab (CBD) for sharing its chemiluminescence imaging system. Some of the images were acquired on Zeiss LSM880 supported by Hercules AKUL/15/37_GOH1816N and FWO G.0929.15 to Pieter Vanden Berghe, KU Leuven. Some of the images were acquired with support from the Zuckerman Institute’s Cellular Imaging Platform. The authors gratefully acknowledge the VIB Bio Imaging Core for their support & assistance in this work. Drawings were created using biorender.com.

## Author contributions

Conceptualization: BLP, PV, FP; Methodology: BLP, RI, PV, FP, KW; Investigation: BLP, AJR, KW, EPH, PV, FP; Formal analysis: BLP, AJR, KW, BV, PV, FP; Critical Reagents: BLP, RI, MD, VG, DR, CC; Funding acquisition: PV, FP; Project administration: PV; Supervision: PV, FP; Writing – original draft: BLP, PV, FP

## Data and materials availability

All data are available in the main text or the supplementary materials. All materials are available from PV.

## Materials and Methods

### Animals

All mouse experiments with xenotransplantation were performed with the approval of the KU Leuven Committee for animal welfare (protocol 2018/030). Mouse housing, breeding and experimental handling were performed according to the ethical guidelines of the Belgian Ministry of Agriculture in agreement with European community Laboratory Animal Care and Use Regulations (86/609/CEE, Journal officiel de l’Union européenne, L358, 18 December 1986). Animals were housed under standard conditions (12 h light:12 h dark cycles) with food and water ad libitum. *Rag2*^-/-^ mice used for xenotransplantations were purchased from Jackson Labs and bred and maintained in local facilities. The day of birth was defined as P0. The xenotransplantations were performed at P0 and animals have been processed between 2 and 18 months.

All mice used to perform *in utero* electroporation (IUE) were handled according to protocols approved by the institutional animal care and use committee (IACUC) at Columbia University. Animals were housed under standard conditions (12 h light:12 h dark cycles) with food and water ad libitum. IUE were performed at embryonic day (E)15.5. The conditional Srgap2 knockout (KO) mouse line was generated as previously described (*1*). The conditional Syngap1 KO mouse line (JAX stock #029303) was obtained from The Jackson Laboratory (*2*). Embryos were staged in days of gestation, where day 0.5 (i.e., E0.5) was defined as noon on the day on which a vaginal plug was detected after overnight mating. C57BL/6J mice were crossed once with the outbred strain 129S2/SvPasCrl mice (obtained from Charles River) to produce F1 hybrid females. Then, timed pregnancies were done by crossing these F1 females with C57BL/6J mice to generate wild-type (WT) animals, with Srgap2^F/F^ mice to generate heterozygous Srgap2^F/+^ animals, and with Syngap1^F/F^ mice to generate heterozygous Syngap^F/+^ animals. To generate the double heterozygous Srgap2^F/+^ ; Syngap^F/+^ animals, timed-pregnant female Srgap2^F/F^ mice were crossed with Syngap1^F/F^ mice.

The data obtained from all animals were pooled without discrimination of sexes for the analysis. Data for this study are derived from a total of 156 mice of both sexes.

### Cell lines and neuronal differentiation

Culture of human pluripotent stem cells (ESC H9) have been described previously (*3*). Human H9 ESC SYNGAP1^+/-^ were generated previously (*4*).

Cells were maintained on irradiated mouse embryonic fibroblasts (MEF) in the ES medium until the start of cortical differentiation. Cortical differentiation from human ESC was performed as described previously (*5*) with some modifications (*6, 7*). On DIV−2, ESCs were dissociated using Stem-Pro Accutase (Thermo Fisher Scientific, Cat#A1110501) and plated on matrigel- (hES qualified matrigel BD, Cat#354277) coated dishes at low confluency (5,000–10,000 cells/cm2) in MEF-conditioned hES medium supplemented with 10 μM ROCK inhibitor (Y-27632; Merck, Cat#688000). On DIV0 of the differentiation, the medium was changed to DDM (*8*), supplemented with B27 devoid of Vitamin A (Thermo Fisher Scientific, Cat#12587010) and 100 ng/ml Noggin (R&D systems, Cat#1967-NG), and the medium was changed every 2 days until DIV6. From DIV6, the medium was changed every day until DIV16. After DIV16, the medium was changed to DDM, supplemented with B27 (DDM/B27), and changed every day. At DIV25, the progenitors were dissociated using Accutase and cryopreserved in mFreSR (StemCell Technologies, Cat#05855).

Differentiated cortical cells were validated for neuronal and cortical markers by immunostaining using antibodies for TUBB3 (BioLegend, Cat#MMS-435P), TBR1 (Abcam, Cat#ab183032), CTIP2 (Abcam, Cat#ab18465), FOXG1 (Takara, Cat#M227), SOX2 (Santa Cruz, Cat#sc-17320), FOXP2 (Abcam, Cat#ab16046), SATB2 (Abcam, Cat#ab34735).

HEK293TT human embryonic kidney cells were obtained from American Type Culture Collection (ATCC cat# CRL-11268). HEK293TT cells were grown in Dulbecco’s modified Eagle’s medium (DMEM; Invitrogen) supplemented with 10% fetal bovine serum (FBS; Invitrogen), 100mM Na-pyruvate, 8.9 mM NaHCO_3_, and penicillin/streptomycin (Invitrogen) and split using TrypLE™ Express Enzyme.

### DNA constructs

pLV-H1-shRNA-hsynapsin-EGFP-WPRE is a modified lentiviral plasmid derived from pH1- SCV2 described in (*9*). The CAG promotor and the mVenus sequences were replaced by hsynapsin and EGFP sequences, respectively, using InFusion cloning (Clontech, Cat#638909). hsynapsin and EGFP sequences were PCR amplified from pLenti-hSynI-EmGFP-WPRE described in (*7*). All constructs were verified by DNA sequencing. The shRNA sequences were inserted by ligation at the NheI/BamhI sites. The shScramble seed sequence was 5’-ACA CCT ATA ACA ACG GTA G-3’ described in (*9*), the shSRAGAP2C#1 sequence was 5’- ACACCTAAAGGTGCAAACATTAA-3’ described in (*9*), the shSRAGAP2C#2 sequence was 5’-AAGGACAGGCATTGAATATCTTA-3’ described in (*9*), the shSRGAP2A sequence was 5’- GCCACTCATCCCTGAAGAATC-3’. All constructs were verified by DNA sequencing.

pLV-hsynapsin-SRGAP2C-HA-WPRE was generated by insertion of SRGAP2C-HA at a multiple cloning site (MCS) of a pLV-hsynapsin-MCS-WPRE plasmid using InFusion cloning (Clontech, Cat#638909). The MCS was prepared by annealing the following primers pair, 5’- CCGGTGCTAGCGTTAACTATATAGGCGCGCCG-3’/5’-

AATTCGGCGCGCCTATATAGTTAACGCTAGCA-3’, inserted into the lentiviral backbone pLenti-hsynapsin-hChR2(H134R)-EYFP-WPRE (a gift from Karl Deisseroth (Addgene plasmid # 20945) by restriction digestion (AgeI/EcoRI) and ligation to obtain a pLV-hsynapsin-MCS-WPRE. SRGAP2C-HA was PCR amplified from the cDNA described in (*9*). All constructs were verified by DNA sequencing.

The pCAG-Cre plasmid was previously described (*10*). The FLEX-TdTomato plasmid was previously described (*11*).

### Virus production

HEK-293T cells were transfected by packaging plasmids, psPAX2 (Addgene Cat#12260) and pMD2.G (Addgene Cat#12259), and the plasmid on interest in a lentiviral backbone. 3 days after transfection, culture medium was collected and viral particles were enriched by filter device (Amicon Ultra-15 Centrifuge Filters, Merck, Cat#UFC910008). Titer check was performed on HEK-293T cell culture for every batch of lentiviral preparation.

### Whole cell protein extraction

Human cortical cells that were frozen at DIV25, were thawed and plated on matrigel-coated plates using DDM/B27 and Neurobasal supplemented with B27 (DDM/B27+Nb/B27) medium. Five days after plating (DIV30), cells were dissociated using Accutase and plated on new matrigel-coated plates at high confluency (100,000–600,000 cells/cm^2^) with lentiviral vector or not infected. The following day, medium was changed to DDM/B27+Nb/B27 medium. 14 days after thawing (DIV39), cells were treated with 10 μM DAPT (Abcam, Cat#ab120633) for 72 hours. The following day (DIV42), cells were treated with 10 μM DAPT and 5 μM Cytarabine (ARA-C) (Merck, Cat#C3350000) for 24 hours. The following day (DIV43), medium was changed to DDM/B27+Nb/B27 medium. At DIV45, cells were homogenized from 2 wells per condition in RIPA buffer (Tris-HCl 50 mM pH = 7.5, NaCl 150mM, NP-40 0,5%, sodium deoxycholate 0,5%, SDS 0.05%, protease inhibitors) using a plastic cell scraper. Samples have been extracted for 1 hours and centrifuged at 10000xg for 30 minutes at 4°C to pellet insoluble material.

Subsequently, samples were diluted to final 1x Laemmli buffer at 95°C for 5mn. They have been run in NUPAGE 10% Bis-Tris Protein Gel at the voltage of 90V for 2 hours in MOPS buffer and then transferred to PVDF Blotting Membrane at the voltage of 100V for 100 minutes. The membrane was blocked in the buffer (5% skim milk and 0.1% Tween20 in TBS) for 1 hour at room temperature and subsequently incubated in the blocking buffer containing rabbit anti- SRGAP2 Nter to label SRGAP2B/C and SRGAP2A (1:1000, described in (*12*) and mouse anti-GAPDH (1:5000, Sigma #G8795). Antibodies were incubated overnight at 4°C, followed by the incubation in the blocking solution containing secondary antibody anti-Rabbit and Mouse IgG antibodies conjugated with HRP at room temperature for 1 hour. Pierce ECL Western Blotting Substrate was used for signal detection.

### Neonatal xenotransplantation

Neonatal xenotransplantation was performed as already described (*5, 7*) with some modifications. Human cortical cells that were frozen at DIV25, were thawed and plated on matrigel-coated plates using DDM/B27 and Neurobasal supplemented with B27 (DDM/B27+Nb/B27) medium. Six days after plating (DIV31), cells were dissociated using Accutase and plated on new matrigel-coated plates at high confluency (100,000–600,000 cells/cm^2^) with lentiviral vector. The following day, medium was changed to DDM/B27+Nb/B27 medium. At 14 days after thawing (DIV39), cells were treated with 10μM DAPT (Abcam, Cat#ab120633) for 72 hours. The following day (DIV42), cells were treated with 10μM DAPT and 5μM Cytarabine (ARA-C) (Merck, Cat#C3350000) for 24 hours. The following day (DIV43), medium was changed to DDM/B27+Nb/B27 medium. 19 days after thawing (DIV44), cells were dissociated using NeuroCult dissociation kit (StemCell technologies, Cat#05715) and suspended in the injection solution containing 20 mM EGTA (Merck, Cat#03777) and 0.1% Fast Green (Merck, Cat#210-M) in PBS at 100 000 cells/μl. Approximately 1-2 μl of cell suspension was injected into the lateral ventricles of each hemisphere of neonatal (P0) immunodeficient mice *Rag2*^−/−^ using glass capillaries pulled on a horizontal puller (Sutter P-97).

### Xenotransplanted mouse cortex processing and immunostaining

Xenotransplanted animals were perfused transcardiacally with ice-cold sucrose 8% PFA 4%. Brains were dissected and soaked in the same fixative overnight, then stored in PBS azide. Then they have been sectioned in 80 μm thickness using vibratome. Slices were transferred into the blocking solution (PBS 0.3% Triton, 5% horse serum, 3% BSA) and incubated for 2 hours. Brain floating slices were incubated 3 days at 4°C with primary antibodies: chicken anti-EGFP (1:1000; ab13970, Abcam), rabbit anti-TBR1 (1:1000, described above), rabbit anti-SATB2 (1:2000, described above), rat anti-CTIP2 (1:500, described above), mouse anti-FOXP2 (1:1000, described above), rat anti-HA (1:500; Roche #11867423001). After three PBS washes, slices were incubated overnight at 4°C with secondary antibodies in PBS: donkey anti-rabbit Cy3, anti-mouse a647, anti-rat a647, anti-chicken a488, anti-rat Cy3 (1:1000) and Hoechst (1:10000). After three washes in PBS, brain sections were mounted on a slide glass with the mounting reagent (DAKO glycerol mounting medium) using #1.5 coverslips.

### Image acquisition of human neurons

Confocal images were obtained with Zeiss LSM880 and LSM900 driven by Zen Black and Blue softwares equipped with objectives 10x, 20x, oil immersion 25x and oil immersion 40x & 63x, AiryScan system and argon, helium-neon and 405 nm diode lasers. Specifically, dendritic branches were acquired using an oil immersion x63 objective and AiryScan system. Neurons immunoreactive for TBR1 and located in cortical layers V-VI of the visual and somatosensory cortices have been considered. For the rescue experiment, EGFP-HA-immunoreactive neurons have been considered. 1-3 proximal dendritic branches from each neuron have been imaged, located 50 µm from the soma.

### Image analysis of human neurons

For human experiments, morphometric analyses of dendritic arbour and dendritic spines were performed by manual 3D reconstruction using Imaris software (Bitplane). The dendritic arbour was reconstructed using the Filament Tracer followed by Sholl analysis. Dendritic spines were quantified in proximal dendrites located 50 µm from the soma. Spine density was defined as the number of quantified spines divided by the length over which the spines were quantified. Head width was defined as the average length of the head that was perpendicular to the neck. The Cell Counter toolbox in Fiji was used to manually annotate cell fate markers.

### Electrophysiological recordings and analysis

Whole cell patch-clamp recordings were performed on acute coronal slices prepared from 6 months old mice with xenotransplanted PSC-derived human neurons. Briefly, animals were anaesthetized intraperitoneally with Nembutal and transcardially perfused with ∼25 mL of ice-cold NMDG-based slicing solution. Brains were rapidly extracted and placed in ice-cold NMDG-based slicing solution containing (in mM): 93 N-Methyl-D-glucamine, 2.5 KCl, 1.2 NaH2PO4, 0.5 CaCl2, 10 MgSO4, 30 NaHCO3, 5 Na-ascorbate, 3 Na-pyruvate, 2 Thiourea, 20 HEPES and 25 D-glucose (pH adjusted to 7.35 with 10 N HCl, gassed with 95% O_2_/5% CO_2_). Coronal slices (250 µm) were cut in ice-cold NMDG-based slicing solution (using a Leica VT1200) and subsequently incubated for ∼6 minutes in the NMDG solution at 34°C. Slices were then transferred into holding aCSF, containing (in mM): 126 NaCl, 3 KCl, 1 NaH2PO4, 1 CaCl2, 6 MgSO4, 26 NaHCO3 and 10 D-glucose (gassed with 95% O_2_/5% CO_2_). Slices were stored at room temperature for ∼1 hour before experiments.

During experiments brain slices were continuously perfused in a submerged chamber (Warner Instruments) at a rate of 3-4 ml/min with 127 mM NaCl, 2.5 mM KCl, 1.25 mM NaH 2 PO 4, 25 mM NaHCO 3, 1 mM MgCl 2, 2 mM CaCl 2, 25 mM glucose at pH 7.4 with 5% CO 2 / 95% O 2 . For sEPSC and AMPA/NMDA ratio recordings we added 20 µM bicuculline. Whole cell patch clamp recordings were done using borosilicate glass recording pipettes (resistance 3.5–5 MΩ, Sutter P-1000) filled with the following internal solution: 115 mM CsMSF, 20 mM CsCl, 10 mM HEPES, 2.5 mM MgCl 2, 4 mM ATP, 0.4 mM GTP, 10 mM Creatine Phosphate and 0.6 mM EGTA. Visually identifiable fluorescently labelled transplanted neurons were selected for recording. Whole-cell patch-clamp recordings were done using a double EPC-10 amplifier under control of Patchmaster v2 x 32 software (HEKA Elektronik, Lambrecht/Pfalz, Germany). Currents were recorded at 20 Hz and low-pass filtered at 3 kHz when stored. The series resistance was compensated to 75-85%. Spontaneous input was recorded using whole-cell voltage clamp recordings (V m =-70 mV). An extracellular stimulation pipette (borosilicate theta glass, Hilgenberg) was placed near the recorded neuron and used for initiating evoked AMPAR and NMDAR-mediated currents (80-120 µA, 1ms (Isoflex, A.M.P. Instruments LTD)). AMPAR-mediated evoked EPSCs were measured in whole-cell voltage clamp at a holding potential of -70 mV, while the NMDAR-mediated component was measured at +40 mV immediately after the initial AMPAR/NMDAR-mediated current (100-150 ms after electrical stimulation). Evoked data were analysed using Fitmaster (HEKA Elektronik, Lambrecht/Pfalz, Germany), spontaneous input was analyzed using Mini Analysis program (Synaptosoft).

### Human synaptosome protein extraction and analysis

Human cortical cells (frozen at DIV25) were thawed and plated on Matrigel-coated plates using DDM/B27+Nb/B27 medium at 37°C with 5% CO_2_. Seven days after thawing (DIV32), the cells were dissociated using Accutase and plated on Matrigel-coated plate at high confluency (450,000- 700,000 cells/cm^2^). Four days later (DIV36), the medium was changed to DDM/B27+Nb/B27 medium with 10µM DAPT. Two days after DAPT induction (DIV38), the medium was changed to fresh DDM/B27+Nb/B27 medium. At DIV40, cortical cells were dissociated using NeuroCult Enzymatic Dissociation Kit following manufacturer’s instructions. For L1CAM+ MACS, dissociated cells were incubated with biotin conjugated anti-human CD171(L1CAM) (Miltenyi Biotec, Cat#130-124-046) in MACS buffer at 4°C for 10 min. After washing using MACS buffer, human cells were incubated with anti-biotin microbeads (Miltenyi Biotec, Cat#130-090-485) in MACS buffer at 4°C for 15 min. L1CAM positive selection were carried out with LS columns according to the manufacturer’s instructions. The sorted cells were plated on Poly-L-ornithine-, Laminin- and horse serum-coated 6-well plate at 110,000 cells/cm^2^. The sorted cells were maintained in DDM/B27+Nb/B27 medium. Neurons have been infected at DIV45 with LV, then the medium was changed to DDM/B27+Nb/B27 medium and changed every 3-4 days until DIV70.

At DIV70, synaptosome proteins have been processed as following. Crude synaptosome extracts were prepared for each condition from 2 wells with 2M neurons, homogenized in Homogenization buffer (sucrose 1.28M, Tris(hydroxymethyl)aminomethane 20mM, MgCl_2_ 4mM, protease inhibitors) using a plastic cell scraper. Homogenate was spun at 1000xg for 10 minutes at 4°C. Supernatant was spun at 14,000 x g for 20 minutes at 4°C. Supernatant was kept as the cytosolic fraction. P2 crude synaptosomes were re-suspended in Extraction Buffer (HEPES 50mM pH = 7.5, NaCl 150mM, EDTA 2mM, NP-40 1%, Triton 0.5%, Na_3_VO_4_ 1mM, NaF 30mM, protease inhibitors) and extracted for 1 hours and centrifuged at 10000xg for 30 minutes at 4°C to pellet insoluble material.

Subsequently, samples were diluted to final 1x Laemmli buffer at 95°C for 5mn. They have been run in NUPAGE 4-12% Bis-Tris Protein Gel at the voltage of 90V for 2 hours in MOPS buffer and then transferred to PVDF Blotting Membrane at the voltage of 100V for 100 minutes. Membranes have been stained using Revert^™^ 700 Total Protein Stain (Licor) following the protocol described by the supplier. The membrane was blocked in the buffer (5% skim milk and 0.1% Tween20 in TBS) for 1 hour at room temperature and subsequently incubated in the blocking buffer containing rabbit anti-SRGAP2 Nter to label SRGAP2C (1:1000, described in (*12*), mouse anti-GAPDH (1:5000, described above), rabbit anti-SYNGAP1 (1:5000; ThermoFisher #PA1- 046), rabbit anti-SRGAP2A (1:5000; Abcam #ab124958), mouse anti-SYP (1:5000; Synaptic Systems #101011), mouse anti-PSD95 (1:5000; Sigma #MABN68) and mouse anti-HOMER1 (1:5000; Synaptic Systems # 160011). Antibodies were incubated overnight at 4°C, followed by the incubation in the blocking solution containing secondary antibody anti-Rabbit and Mouse IgG antibody conjugated with HRP at room temperature for 1 hour. SuperSignal™ Western Blot Substrate were used for signal detection.

Signals have been measured using Fiji and normalized to the total protein staining and then to the WT shScramble value.

### In Utero Electroporation (IUE)

IUE was performed at embryonic day 15.5 (E15.5) on isoflurane-anaesthetized timed-pregnant female mice as previously described (*13*), but with the following modifications. Endotoxin-free DNA containing 1 ug/ul of FLEX-TdTomato plasmid and 2-100 ng/ul of pCAG-Cre plasmid was injected into the ventricles of E15.5 embryos using a heat-pulled capillary attached to Picospritzer III (Parker). Electroporation was performed by applying 5 pulses of 42 V for 50 ms with 500-ms intervals using a 3-mm diameter platinum tweezer electrode (Nepa Gene) and a square wave electroporator (ECM 830, BTX). After placing embryos back into the abdominal cavity, the incision was closed using sutures and the mouse was allowed to recover on a heating pad.

### IUE mouse processing and dendritic spine analysis

At P20-P21, mice were anesthetized with isoflurane, and intracardiac perfusion was performed using 4% paraformaldehyde (Electron Microscopy Sciences, cat no. 15714-S) in PBS as previously described (*14*), but with the following modifications. Brains were isolated and incubated overnight in the 4% paraformaldehyde in PBS solution at 4°C. The brains were then washed in PBS and sectioned along the coronal plane at 100 µm using a vibrating microtome (Leica VT1200S). Sections were collected spanning the approximate bounds of the somatosensory cortex. To confirm L2/3 cortical targeting, representative WT sections were blocked in 10% goat serum and 0.5% Triton X-100 in PBS for 1 hour at room temperature, and then incubated overnight at room temperature with the following antibody: rabbit anti-CUX1 (1:1000; Proteintech, cat no. 11733-1-AP) in 2% goat serum and 0.5% Triton X-100 in PBS. The stained slices were then washed in PBS with 0.5% Triton X-100 and incubated with the following secondary antibody: goat anti-rabbit Alexa Fluor 488 (1:200; Invitrogen, cat no. 11008) in 2% goat serum and 0.5% Triton X-100 in PBS. Sections were stained using DAPI (Invitrogen, cat no. D1306) and subsequently mounted on glass slides in Citifluor Mountant Solution (Electron Microscopy Sciences, cat no. AF100-5 & 17977-150). Slides were imaged on a W1-Tokogawa spinning disk confocal microscope using a Nikon 100x, 1.35 NA silicon-immersion objective (Nikon, cat no. MRD73950).

Morphometric analyses of dendritic spines were performed in the depth of the z stack for slices using Fiji as previously described (*9*), but with the following modifications. Dendritic spines were quantified in oblique dendrites originating from the apical trunk. Head width was defined as the largest length of the head that was perpendicular to the neck. Only dendrites that were parallel to the plane of the slice were analyzed. Spine analysis was done in brain sections of comparable rostro-caudal position.

### Statistical analysis

Data are represented as mean + standard error of the mean (SEM) if no precision or +/- SEM or +/- 95% confidence interval (see Figures for details). Unpaired Mann-Whitney test have been used for assessing the significance of differences in the analyses between two conditions using GraphPad. Kolmogorov-Smirnov test have been used for assessing the significance of differences in the analyses of cumulative distributions using GraphPad. ANOVA tests have been used in the analyses containing more than two conditions using GraphPad (Kruskal-Wallis tests with Dunn’s correction, 2-way ANOVA multiple comparison or 2-way ANOVA tests). See Figures for details. Numbers of litters, animals and neurons used are described in Figure legends and specifically for the Figure 2A-C in Table 1. For xenotransplantation experiments with wild-type neurons, experimental groups were composed of neurons processed and transplanted in the same time, meaning animals were littermates. For the xenotransplantation experiments with SYNGAP1^+/-^ neurons, SYNGAP1^+/-^ neurons with LV-shScramble and LV-shSRGAP2A infection were processed and transplanted in the same time, meaning animals were littermates, but wild-type neurons taken as a comparison were originated from other batches of xenotransplantations. For human synaptosome blots, 3-5 protein extractions have been performed per condition from independent differentiation batches.

---

**Fig. S1.**
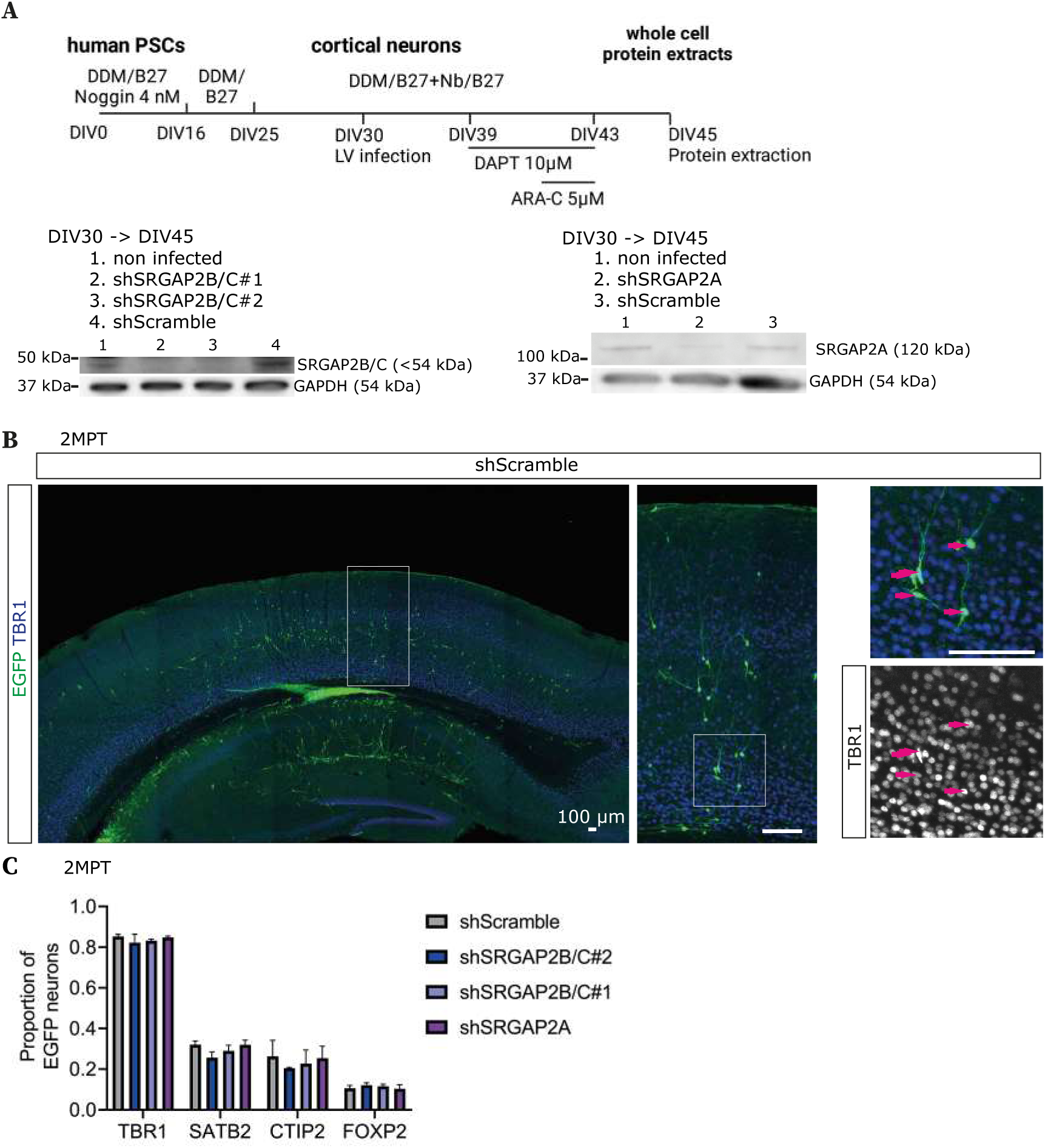
Generating a model of SRGAP2 knock-down (KD) in human cortical pyramidal neurons in vivo. (**A**) Experimental design for the shRNAs validation: pluripotent stem cell (PSC)- derived human neurons were infected with lentivirus (LV) expressing shRNAs at day in vitro (DIV) 30 and whole cell protein extraction was performed at DIV45. Immunoblot for SRGAP2 proteins and GAPDH from whole cell protein extracts of human PSC-derived neural cultures at DIV45 after LV infections at DIV30 (see Fig. S3 for quantification of the shRNAs efficiency). (**B**) Representative picture of a mouse cortical sections with 2MPT human neurons LV-shScramble infected and immunostained for EGFP and TBR1; arrows indicate TBR1 positive EGFP neurons. (C) Quantification of the proportion of 2MPT human neurons LV-shRNA infected and immunoreactive for TBR1, CTIP2, SATB2 and FOXP2 (per condition, 131-245 EGFP neurons from 2 animals from 1 litter). Data are represented as mean + SEM.

**Fig. S2.**
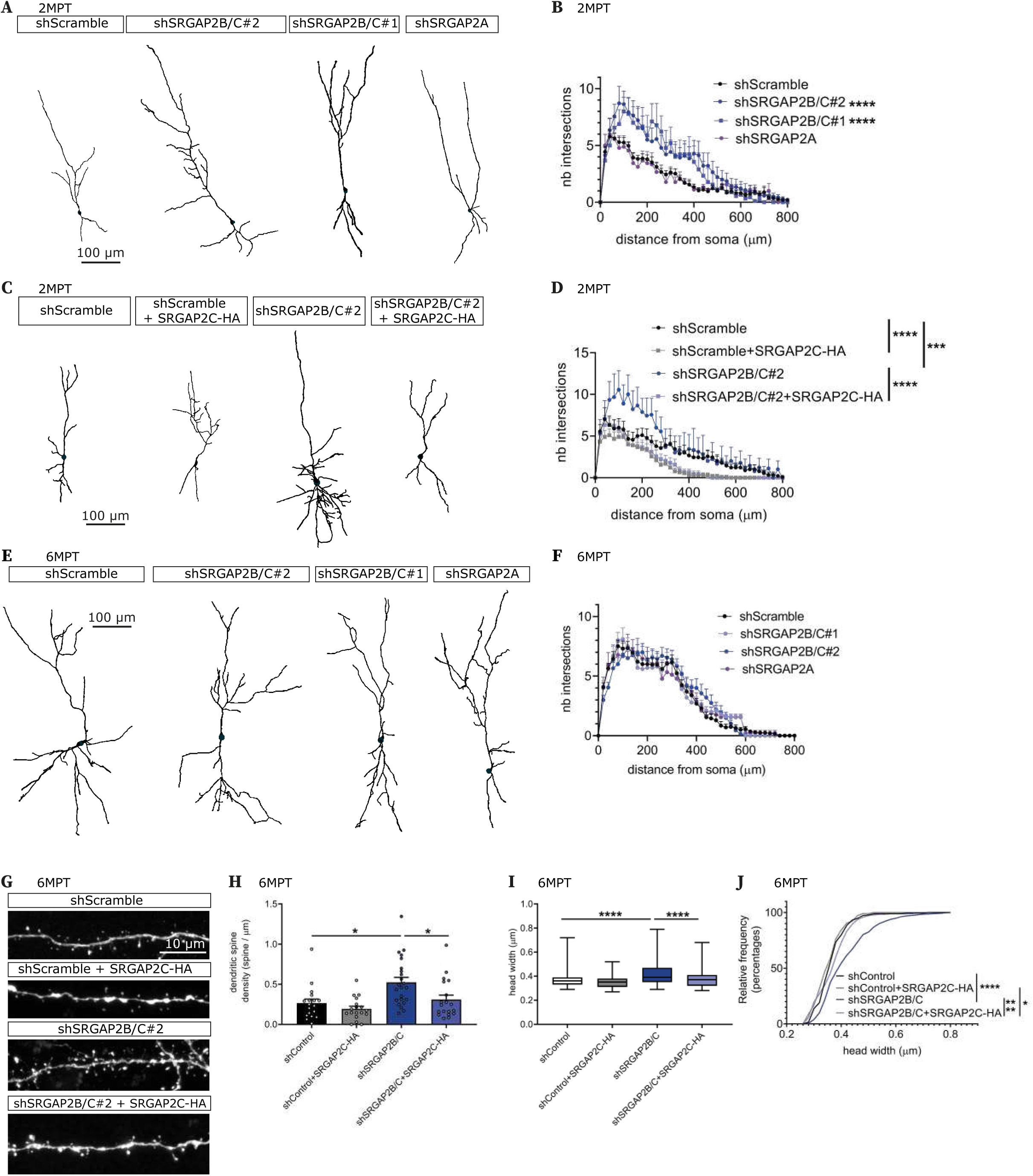
SRGAP2B/C genes are required for human neuronal maturation neoteny. (**A-B**), 3D reconstruction of the dendritic arbour of 2-month post-transplantation (MPT) neurons with LV-shRNA infection and corresponding quantification (2-way ANOVA tests)(shScramble: 27 neurons from 4 litters, 9 animals; shSRGAP2B/C#2: 14 neurons from 2 litters, 5 animals; shSRGAP2B/C#1&shSRGAP2A: 9 neurons from 1 litter, 3 animals, each). (**C-D**), Reconstruction of the dendritic arbour of 2MPT neurons with LV-shRNA +/- SRGAP2C-HA infection and corresponding quantification (2-way ANOVA tests)(9 neurons from 3 animals from 1 litter for each condition). (**E-F**) 3D reconstruction of the dendritic arbour of 6MPT human neurons with LV-shRNA infection and corresponding quantification (2-way ANOVA tests)(shScramble: 20 neurons from 4 litters, 9 animals; shSRGAP2B/C#2: 20 neurons from 2 litters, 6 animals; shSRGAP2#1&shSRGAP2A: 9 neurons from 1 litter, 3 animals, each). (**G**) Representative proximal dendritic branches of human PSC-derived neurons xenotransplanted in the mouse cerebral cortex, with LV-shRNA +/- SRGAP2C-HA cDNA infection at 6MPT. (**H-I**), Corresponding quantifications of the dendritic spine density and spine head width (Kruskal-Wallis tests)(18-22 neurons, 228-774 dendritic spines from 6 animals from 2 litters per condition). (J) Cumulative distribution of the spine head width (Kolmogorov-Smirnov tests). In B, D, F, H, data are represented as mean+SEM. *, p<0.05; ***, p<0.001; ****, p<0.0001.

**Fig. S3.**
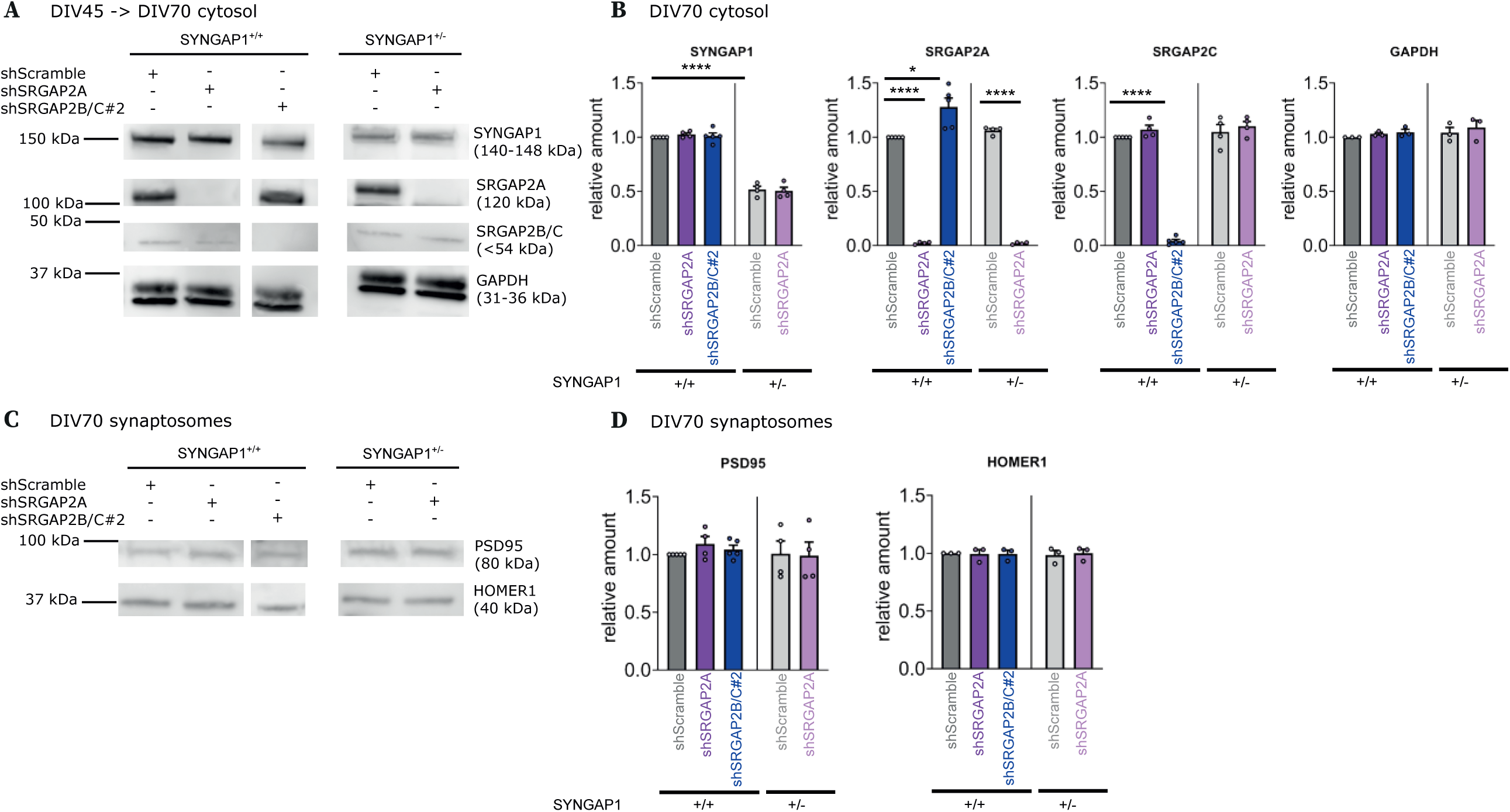
Cytosolic and synaptic fractions of human neurons at DIV70. (**A**) Complementary data to Fig. 3: cytosolic fraction of SYNGAP1+/+ and SYNGAP1+/- PSC-derived neurons at DIV70 with LV-shRNA infection at DIV45 stained for SYNGAP1, SRGAP2A, SRGAP2B/C and GAPDH. (**B**) Corresponding quantification (ANOVA multiple tests; 3-5 experiments per condition). (**C**) Complementary data to Fig. 3B-C: synaptic fraction of SYNGAP1+/+ and SYNGAP1+/- PSC-derived neurons at DIV70 with LV-shRNA infection at DIV45 stained for PSD95 and Homer1. (**D**) Corresponding quantifications (ANOVA multiple tests; 3-5 experiments per condition). Dara are represented as mean + SEM. *, p<0.05. ****, p<0.0001.

**Table S1.** Numbers of litters, animals, neurons and dendritic spines of human xenotransplantated neurons (related to Figure 1C-E)

|  | n litters | n animals | n neurons | n branches | n spines |
| --- | --- | --- | --- | --- | --- |
| 2MPT |  |  |  |  |  |
| shSRGAP2B/C#2 | 3 | 6 | 30 | 45 | 226 |
| shSRGAP2B/C#1 | 1 | 2 | 10 | 15 | 101 |
| shSRGAP2A | 1 | 2 | 10 | 10 | 65 |
| shScramble | 4 | 8 | 40 | 48 | 190 |
| 4MPT |  |  |  |  |  |
| shSRGAP2B/C#2 | 3 | 6 | 27 | 66 | 1254 |
| shScramble | 3 | 6 | 27 | 56 | 339 |
| 6MPT |  |  |  |  |  |
| shSRGAP2B/C#2 | 3 | 6 | 30 | 60 | 1970 |
| shSRGAP2B/C#1 | 3 | 8 | 28 | 56 | 1690 |
| shSRGAP2A | 2 | 5 | 15 | 30 | 268 |
| shScramble | 8 | 14 | 72 | 123 | 1440 |
| 9MPT |  |  |  |  |  |
| shSRGAP2B/C#2 | 1 | 3 | 18 | 37 | 1959 |
| shScramble | 1 | 2 | 19 | 37 | 753 |
| 12MPT |  |  |  |  |  |
| shSRGAP2B/C#2 | 2 | 6 | 19 | 38 | 2005 |
| shSRGAP2B/C#1 | 2 | 5 | 20 | 38 | 1686 |
| shSRGAP2A | 2 | 4 | 12 | 24 | 261 |
| shScramble | 6 | 2 | 33 | 65 | 1808 |
| 18MPT |  |  |  |  |  |
| shSRGAP2B/C#2 | 1 | 2 | 9 | 18 | 2007 |
| shSRGAP2B/C#1 | 1 | 2 | 9 | 18 | 922 |
| shSRGAP2A | 1 | 2 | 9 | 18 | 300 |
| shScramble | 1 | 2 | 9 | 18 | 775 |

